# PXN/Paxillin Phase Separation Promotes Focal Adhesion Assembly and Integrin Signaling

**DOI:** 10.1101/2022.12.17.520852

**Authors:** Peigang Liang, Yuchen Wu, Shanyuan Zheng, Jiaqi Zhang, Shuo Yang, Jinfang Wang, Suibin Ma, Mengjun Zhang, Zhuang Gu, Qingfeng Liu, Wenxue Jiang, Qiong Xing, Bo Wang

## Abstract

Focal adhesions (FAs) are transmembrane protein assemblies mediating cell-matrix connection. Tools to manipulate the compositionally intricate and dynamic FAs are currently limited, rendering many fundamental hypotheses untestable. Although protein liquid-liquid phase separation (LLPS) has been tied to the organization and dynamics of FAs, the underlying mechanisms remain unclear. Here, we experimentally tune the LLPS of PXN/Paxillin, an essential scaffold protein of FAs, by utilizing light-inducible Cry2 system. In addition to nucleating FA components, light-triggered PXN LLPS potently activates integrin signaling and subsequently accelerates cell spreading. PXN favors homotypic interaction-driven LLPS *in vitro*. In cells, PXN condensates are associated with plasma membrane, and modulated by actomyosin contraction and client proteins of FAs. Interestingly, non-specific weak inter-molecular interactions, together with specific molecular interactions, underlie the multicomponent condensation of PXN, and are efficient to promote FA assembly and integrin signaling. Thus, our data establish an active role of PXN phase transition into a condensed membrane-associated compartment in promoting assembly/maturation of FAs.

## Introduction

Liquid-liquid phase separation (LLPS) is a biophysical phenomenon when proteins spontaneously demix in solutions (Alberti et al., 2019; Alberti and Hyman, 2021). Even since its first linkage to P-granule dynamics in the *C.elegans*, LLPS has now been implicated in a myriad of biological activities (Brangwynne et al., 2009). It is well-accepted that LLPS underlies the formation of membrane-less compartments, such as nucleolus and stress granules (Shin and Brangwynne, 2017). Aberrant LLPS leads to disturbed dynamics of membrane-less organelles, which is thought to be the main cause of many diseases, especially neurodegeneration and cancer (Alberti et al., 2019; Alberti and Hyman, 2021; Mathieu et al., 2020). Proteins with LLPS properties often harbor unstructured regions, also known as intrinsically disordered regions (IDRs), which render the proteins readily for either intra- or inter-molecular weak promiscuous interactions (Feng et al., 2021). In addition, protein interaction motifs promote LLPS and provide “multivalency” to the membrane-less compartments by interacting with binding partners stoichiometrically (Feng et al., 2021). Thus far, the molecular grammar governing LLPS has been mostly elucidated in well-studied membrane-less systems, such as stress granules. It is likely that diverse mechanisms exist to regulate the properties and functions of other biological condensates under various pathophysiological conditions.

One of the most remarkable applications of LLPS is in the biosynthesis of artificial membrane-less organelles (Bracha et al., 2019). By fusing protein fragments that drive LLPS to protein elements that are capable of undergoing light-, temperature-, or chemical-dependent homotypic or heterotypic oligomerization, controllable condensation is achieved spatiotemporally (Garabedian et al., 2021; Schuster et al., 2018). Importantly, these de novo synthetic organelles often exhibit desired biophysical and biochemical properties (Garabedian et al., 2021; Schuster et al., 2018; Zhao et al., 2019). The tunable features of synthetic organelles permit precise delineation of signaling pathways and regulatory mechanisms underlying diverse cell behaviors, which otherwise remain challenging for traditional biochemical or molecular approaches. For example, persistent stress granules are long believed to be cytotoxic and contribute to the pathogenesis of several neurodegenerative diseases, such as amyotrophic lateral sclerosis (Li et al., 2013). This is only conceptually well-accepted but unprovable because of the technical hurdles to generate persistent stress granules *in vitro*. Utilizing the LLPS of core stress granule protein G3BP1, persistent designer stress granules are produced by blue light and shown to cause marked cytotoxicity for the first time (Zhang et al., 2019). Despite the appealing future of synthetic organelles, it has only been successfully applied to the prokaryotic cells or a limited number of organelles (Garabedian et al., 2021; Zhang et al., 2019). Inventions of more versatile synthetic organelles will be valuable tools for biomedical applications and beyond.

FAs are transmembrane structures that connect the extracellular matrix and cellular cytoskeleton to govern tissue morphogenesis, mechanosensing, and cell migration. The functions of FAs are particularly reliant on integrins, which are α/β heterodimeric transmembrane receptors binding to the extracellular ligands or the receptors of neighboring cells. Integrins relay chemical or mechanical cues to intracellular cytoskeletal systems by recruiting various scaffold proteins through the cytoplasmic tails (Winograd-Katz et al., 2014). FAs constantly evolve in size, shape, constitution, and hence property in response to different extracellular and intracellular cues. The lifecycle of FAs starts with the earliest adhesions forming at the cell front, known as nascent adhesions, which then either collapse or develop into mature adhesions, termed FAs. During this process, extensive morphological and compositional remodeling occur (Parsons et al., 2010). Unbiased proteomic approaches have identified hundreds of proteins in the FAs (Horton et al., 2015; Kuo et al., 2011; Robertson et al., 2015). These protein machineries participate in various heterotypic and/or homotypic protein-protein interactions via diverse modular domains (Geiger and Yamada, 2011; Horton et al., 2015). Much of the efforts in the last several decades have been devoted to uncover the molecular and biochemical mechanisms through which the adhesions arise, mature and disassemble, with particular focus on the protein-protein interactions (Geiger and Yamada, 2011; Parsons et al., 2010). Nevertheless, the sheer complexity of the organizing principles and regulatory mechanisms of FAs confound our understanding and impede attempts to reconstitute adhesions.

Integrin orchestrates high order assembly of hundreds of proteins through the cytoplasmic tail. Therefore, this cytoplasmic macromolecular complex of FAs is membranes-less. Apart from the membrane-less and dynamic features, the cytoplasmic assembly of FAs also exhibits liquid-like properties, including constant fusion and material exchange with the surroundings. We hypothesis that LLPS of FA associated-proteins contributes to the assembly of the cytoplasmic compartment of FAs. Only until recently, few studies have demonstrated that proteins in the FAs including GIT1, LIMD1, FAK, and Kindlin-2 undergo LLPS (Case et al., 2022; Li et al., 2020; Wang et al., 2021; Zhu et al., 2020), albeit it is less clear whether these proteins regulate adhesion dynamics through LLPS. Here, by manipulating the LLPS of PXN, a key scaffold protein of FAs, using previously developed optogenetic tools (Bracha et al., 2018; Shin et al., 2017), we successfully synthesize de novo membrane-associated macromolecular assemblies, which efficiently concentrate cargo proteins belonging to the FA proteome, and faithfully recapitulate key molecular events during FA formation and maturation. More importantly, light-induced synthetic adhesions are sufficient to activate integrin signaling and facilitate cell spreading. Further, we find that *in vitro*, PXN undergoes phase separation initiated by homotypic interactions, but is prone to form complex condensation when other client proteins are present under physiological conditions and that the material properties of PXN condensates are modulated by actomyosin contraction. Interestingly, non-specific protein-protein interactions plays an under-appreciated, yet essential role in the complex condensation of PXN. Together, we propose that PXN phase separation, along with other well-established mechanisms such as protein-liquid interactions, retrograde actin flow, and actomyosin contraction, contribute to FA assembly and maturation.

## Results

### Tuning PXN LLPS in cells using an optogenetic approach

Inspired by previously developed optical system where disordered regions of specific proteins are fused with Cry2_PHR_-mCherry, which undergoes blue light-dependent oligomerization, providing the initial trigger for condensation (Shin et al., 2017). This approach successfully achieved designed stress granules (Zhang et al., 2019). To test whether optically controlled LLPS of a scaffold protein of FAs could faithfully reconstitute a FA-like structures, we chose PXN/Paxillin because our own serendipitous results showed that PXN was required for FA formation specifically in HeLa cells (Fig. S1 A). PXN consists of 5 LD domains located at the N-terminus participating in extensive protein-protein interactions and 4 LIM domains aligned in tandem at the C-terminus involved in binding to additional proteins (Alpha et al., 2020; Deakin and Turner, 2008). The N-terminus was predicted to be mostly disordered, we therefore fused this fragment with Cry2_PHR_-mCherry. We referred to this fusion protein as PXN-Cry2 and mCherry-Cry2 alone as Control-Cry2 (Fig. 1 A). Expression of Control-Cry2 or PXN-Cry2 had no appreciable impact on the disrupted FA formation in the *PXN*^-/-^ HeLa cells. While *PXN*^-/-^ HeLa cells stably expressing Control-Cry2 remained unresponsive to brief blue light exposure, cells with PXN-Cry2 instantly formed numerous condensates upon blue light treatment, which were named as Opto-PXN (Fig. 1 B). The Opto-PXN droplets spontaneously disassembled within 10 minutes after the blue light treatment ceased (Fig. 1 C). The Opto-PXN droplets could fuse (Fig. 1 D) and exchange materials with the surrounding cytoplasm (Fig. 1, E and F). Additionally, the Opto-PXN underwent cycles of formation and dissolution (Fig. S1 B), and the formation was profoundly inhibited by 1,6-hexanediol treatment (Fig. S1 C), suggesting contributions from hydrophobic forces to the light-elicited PXN LLPS. HIC-5 is a close paralogue of PXN sharing extensive sequence and structural similarities (Deakin and Turner, 2008; Thomas et al., 1999). Both redundant and non-redundant functions have been reported for PXN and HIC-5 (Deakin and Turner, 2011). we asked if HIC-5 is capable of forming droplets under similar experimental conditions. Intriguingly, HIC-5-Cry2 failed to produce condensates (Fig. 1, G and H) despite that HIC-5-Cry2 expression was similar to that of PXN-Cry2 (Fig. S1 D). Given that LLPS is highly dependent on the protein concentration, we therefore hypothesized that the induction of Opto-PXN assembly would be dependent on the local concentration of PXN. We controlled the local PXN concentration by modulating either the intensity of the activating blue light or the expression level of the PXN-Cry2 fusion protein (indicated by the relative fluorescence intensity of mCherry). As predicted, we observed a strong positive correlation between blue light intensity and induction of Opto-PXN (Fig. 1, I and J). Besides, a strong positive correlation between PXN-Cry2-mCherry expression level and induction of Opto-PXN was also established (Fig. 1, K and L).

**Figure 1.**
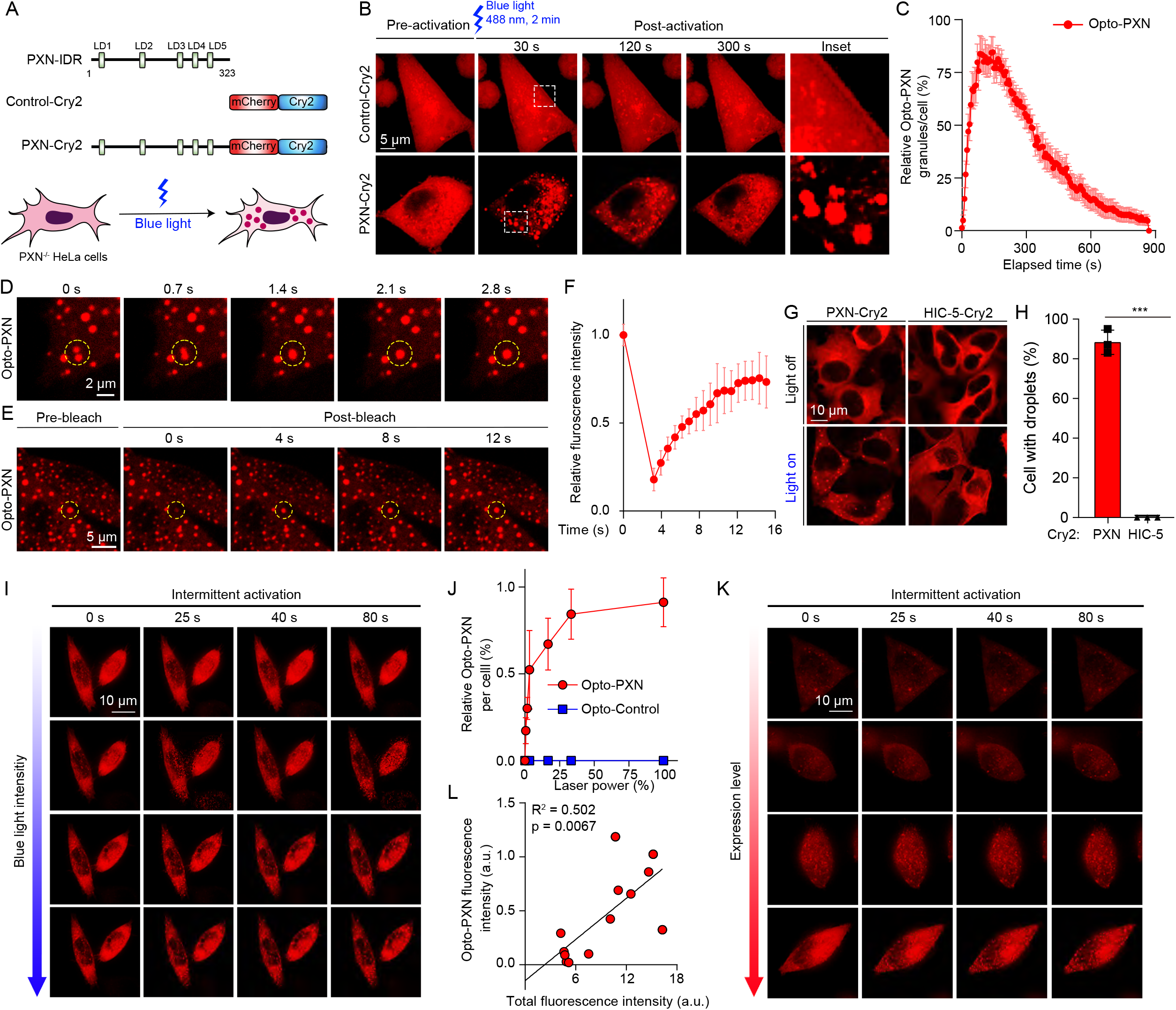
Light-induced PXN LLPS. (**A**) Schematic illustrations of the Control-Cry2 and PXN-Cry2. (**B**) Representative images of *PXN*^-/-^ HeLa cells stably expressing Control-Cry2 or PXN-Cry2 constructs treated with 2 min of 488 nm blue light. (**C**) Quantification of the Opto-PXN droplet formation over time from (**B**). Data are shown as mean ± SD. (**D**) Live images show the rapid fusion of Opto-PXN. (**E**, **F**) HeLa cells expressing PXN-Cry2 were first stimulated with a pulse of 488 blue light, and the circled areas were photobleached. The fluorescence intensity was monitored over time and quantified (**F**). Data are shown as mean ± SD. n = 9. (**G**) Representative images of WT HeLa cells stably expressing PXN-Cry2 or HIC-5-Cry2 stimulated with blue light for 30 min. Percentage of cells with droplets were calculated from 3 independent experiments (**H**). Data are shown as mean ± SEM. ***p < 0.001 by unpaired Student t-test. (**I**) HeLa cells expressing PXN-Cry2 were treated with intermittent blue light laser for the indicated time. Blue light intensity was gradually augmented from top to bottom. The correlation between relative blue light laser power and formation of Opto-PXN condensates was determined in (**J**). Data are shown as mean ± SD. (**K**) Cells with different levels of PXN-Cry2 were treated with intermittent blue light for the indicated time followed by image acquisition using the 561 nm channel. The correlation between the relative PXN-Cry2 protein level and the total fluorescence intensity of Opto-PXN condensates was determined in (**L**). Data are shown as mean ± SD.

### Light-induced PXN LLPS drives the selective compartmentation of FA proteins

To define the relationship between the light-induced Opto-PXN condensates and FAs, we next examined their composition. We asked whether PXN recruits FA proteins through light-triggered LLPS. We found that with 10 minutes of blue light illumination, the Opto-PXN droplets efficiently nucleated endogenous FA-associated proteins including HIC-5, FAK, Vinculin, Talin-1, GIT1, and Kindlin-2 (Fig. 2, A and B; and Fig. S1, E and F). Interestingly, many of these proteins, such as FAK and Vinculin, are reported to be present in the early adhesions (Geiger and Yamada, 2011; Parsons et al., 2010). However, components that are excluded in the early adhesions but enriched in the mature adhesions, most of which are F-actin crosslinking proteins including LIMD1, α-actinin, TSN3, VASP, and Zyxin, failed to be detected in the Opto-PXN droplets at 10 min post blue light treatment (Fig. 2, C and D; and Fig. S1, E and F). With extended blue light exposure (30-60 min) however, the Opto-PXN granules gradually recruited the F-actin crosslinking proteins and started to exhibited compositional features of mature adhesions (Fig. 2, C and D; and Fig. S1, E and F). We speculate that the time of appearance and degree of partition of these adhesion proteins in the Opto-PXN droplets (Fig. 2 E) are reflections of the molecular cascade occurring during bona fide FA assembly. It also reminds us with the stratified model of the FAs with PXN and FAK in the membrane-apposed integrin signaling layer, Talin-1 and Vinculin in the intermediate force-transduction layer, and α-actinin, VASP, and Zyxin in the uppermost actin-regulatory layer (Kanchanawong et al., 2010). Although the Opto-PXN droplets are liquid-like and exhibit no polarized distribution of client proteins within the droplets, we suspect other factors under physiological settings such as actin remodeling during FA maturation, which was not modelled by the light-induced PXN condensates, could induce this molecular rearrangement. In line with this conjecture, F-actin was not enriched in the Opto-PXN condensates (Fig. 2 F). Finally, we asked if the Opto-PXN condensates contained integrins. Indeed, the Opto-PXN granules were colocalized with integrin ß1 with prolonged blue light treatment (Fig. 2, G and H, and Fig. S1, E and F), suggesting the synthetic Opto-PXN condensates were compositionally similar to the FAs. Because FAs are plasma membrane-associated cellular compartments, we employed total internal reflection fluorescence (TIRF) microscopy to determine the cellular location of Opto-PXN condensates relative to the plasma membrane. We found that the Opto-PXN condensates were in close proximity to the plasma membrane and gradually grew in size (Fig. 2, I and J). In approximately 50% of the cells, all the Opto-PXN condensates were visible by TIRF (Fig. 2, K and L). Further, we performed plasma membrane fractionation experiment to determine if the Opto-PXN condensates were plasma membrane-associated. Indeed, the synthetic Opto-PXN condensates were associated with plasma membrane, which increased over time with blue light-induced PXN LLPS (Fig. 2 M). Consistently, representative proteins of cytosolic membrane-less compartments, including G3BP1 of the stress granules and YBX1 of the P-body were excluded from the Opto-PXN condensates (Fig. S1, G-J). Conversely, we also generated Opto-FUS, a well-studied light-inducible condensate utilizing the IDR of RNA binding protein FUS. As expected, blue light treatment robustly elicited cytoplasmic condensates. Nevertheless, the Opto-FUS droplets were not immunopositive for PXN or HIC-5 (Fig. S1, K-N). Since the above-described observation were made in the *PXN*^-/-^ HeLa cells, we also stably expressed PXN-Cry2 in the WT HeLa cells. We affirmed that the Opto-PXN condensate markedly enriched FA-resident proteins even in the presence of pre-existing FAs (Fig. S1 O). Collectively, our data suggest that the membrane-attached synthetic adhesions selectively compartmentalize cargo proteins via PXN LLPS.

**Figure 2.**
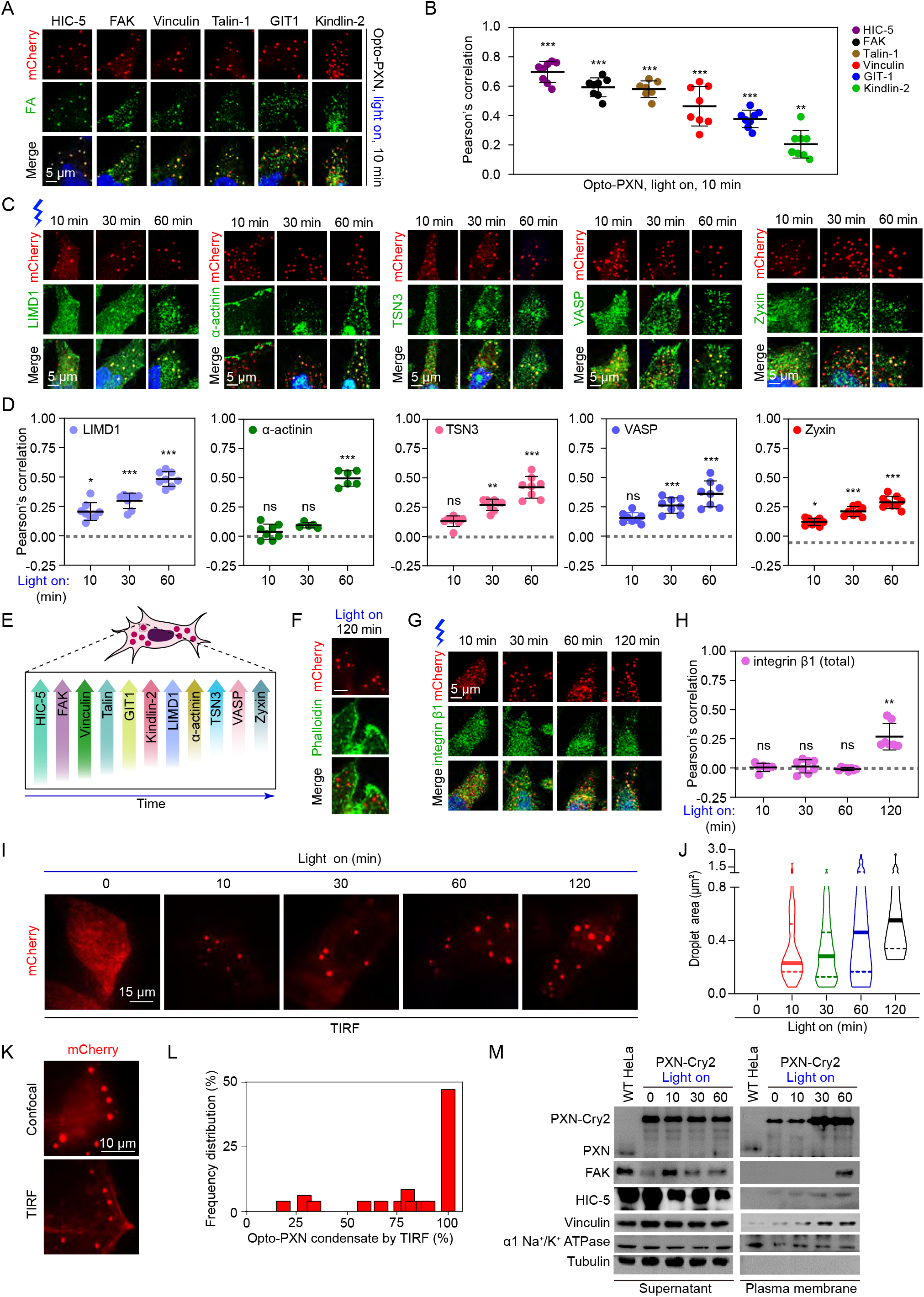
Light-induced PXN LLPS drives the nucleation of FA proteins. (**A**) *PXN*^-/-^ HeLa cells expressing PXN-Cry2 were activated with blue light for 10 min and processed for immunostaining with the indicated antibodies. The Pearson’s correlation of each protein with the Opto-PXN droplets was quantified (**B**). Data are presented as mean ± SD. (**C**-**D**) *PXN*^-/-^ HeLa cells expressing PXN-Cry2 were activated with blue light for the indicated time, and immunostained with antibodies against the indicated proteins (**C**). The Pearson’s correlation was quantified in (**D**). Data are presented as mean ± SD. (**E**) Schematic summary of the results from (**A**-**D**). (**F**) *PXN*^-/-^ HeLa cells expressing PXN-Cry2 were treated with blue light for 120 min and stained with Phalloidin to visualize F-actin. (**G**-**H**) *PXN*^-/-^ HeLa cells expressing PXN-Cry2 were activated with blue light for the indicated time, and immunostained with antibodies against integrin β1 (**G**). The Pearson’s correlation was quantified in (**H**). (**I**-**J**) *PXN*^-/-^ HeLa cells expressing PXN-Cry2 were activated with blue light for the indicated time and were imaged with TIRF microscopy. Droplet area was determined and shown in (**J**). (**K**-**L**) *PXN*^-/-^ HeLa cells expressing PXN-Cry2 were activated with blue light and imaged by confocal and TIRF microscopy, respectively. The percentage of Opto-PXN condensates visible by TIRF was determined (**L**). n = 44 cells. (**M**) *PXN*^-/-^ HeLa cells expressing PXN-Cry2 were activated with blue light for the indicated time and then subjected to plasma membrane fractionation. The samples were analyzed by immunoblotting with the indicated antibodies. ns, not significant; *p < 0.05; **p < 0.01; ***p < 0.001 by one-way ANOVA.

### Light-induced PXN LLPS potentiates integrin signaling and accelerates cell spreading

FAs not only physically connect the extracellular matrix and intracellular cytoskeleton, but are also signaling hubs. Having established the morphological and molecular signature of the light-induced PXN assemblies, we asked if Opto-PXN droplets are capable of transducing integrin-associated signaling. Downstream signaling pathways of integrin include activation of the FAK, Src, AKT, ERK, and JNK pathways and regulation of small GTPases of the RHO family (Legate et al., 2009), which are essential for many integrin-dependent processes including regulation of cytoskeletal rearrangement and cell migration. We next collectively examined the activities of these pathways. Surprisingly, we observed a striking increase in the phosphorylation level of Src at Tyr416 in the blue light treated Opto-PXN expressing cells in a time-dependent fashion, indicating the activation of Src kinase (Fig. 3 A, and Fig. S2 A). Meanwhile, inhibitory phosphorylation of Src at Tyr529 remained unchanged (Fig. 3 A, and Fig. S2 A). The full activation of FAK is regulated by phosphorylation at multiple tyrosine residues (Mitra et al., 2005). Among these, both the autophosphorylation at Tyr397 and phosphorylation at Tyr925 by Src were significantly enhanced by Opto-PXN droplet formation (Fig. 3 A, and Fig. S2 A). As a result of activated FAK and/or Src kinases, phosphorylation of PXN at Tyr31 and Tyr118 also displayed similar increasing trends over time (Fig. 3 A, and Fig. S2 A). More importantly, immunostaining showed that the enhanced phosphorylation of PXN at Tyr31, Tyr118, and FAK at Tyr397 occurred within the Opto-PXN droplets (Fig. 3, B-G). Likewise, AKT, ERK, and JNK were all activated since increased phosphorylation of AKT at Ser473, ERK at Thr202/Tyr204, and JNK at Thr183/Tyr185 were detected in the PXN-Cry2 expressing cells upon blue light exposure (Fig. 3 A, and Fig. S2 A). These data implicate the activation of integrin signaling. To consolidate these findings, we took advantage of a well-characterized antibody (12G10) that specifically recognizes the active integrin ß1 receptor. As expected, immunostaining revealed selective partitioning of activated integrin ß1 into the Opto-PXN droplets (Fig. 3, H and I). Importantly, flow cytometry showed that the overall cellular integrin activity was also drastically enhanced with 60 min or longed blue light treatment in cells harboring PXN-Cry2 but not Control-Cry2 (Fig. 3 J). In contrast, inactive integrin ß1 (recognized by mAb13 antibody) was not found in the Opto-PXN condensates (Fig. S2, B and C). To determine if the phenomenon of Opto-PXN was cell type specific, we generated *Pxn*^-/-^ mouse embryonic fibroblasts (MEFs), wherein the FA assembly/maturation was not entirely blocked (Hagel et al., 2002) and stably reconstituted Cry2-PXN into these cells. Opto-PXN condensates were successfully synthesized with blue light illumination, which markedly recruited FA-resident proteins including FAK, HIC-5, and Vinculin, and clustered active integrin ß1 receptors (Fig. S2 D). Notable upregulation of FAK and PXN phosphorylation was also achieved with blue light-induced PXN LLPS in the MEFs (Fig. S2 E). Finally, we sought to determine if the activated integrin signaling resulted from forced PXN LLPS was functionally relevant. Integrin signaling activation in the short-term facilitates cell attachment and spreading, and in the long-term regulates cell migration (Geiger and Yamada, 2011). Considering the background toxicity imposed by prolonged light treatment (several hours), we decided to focus on the short-term (within 120 min) cell adhesion and spreading. We found that light-induced synthesis of Opto-PXN droplets significantly accelerated cell spreading on the fibronectin-coated surface (Fig. 3, K and L), whereas light treatment had minimal impact on the spreading of the Control-Cry2 expressing cells (Fig. S2, F and G). Note that the accelerated cell spreading only occurred after 60 mins of blue light stimulation, a perquisite for full activation of integrin signaling in our experimental system. Altogether, light-induced LLPS of PXN potently drives condensation of client proteins, and accelerates biochemical reactions in the integrin pathway, and hence facilitates cell spreading (Figure 3M).

**Figure 3.**
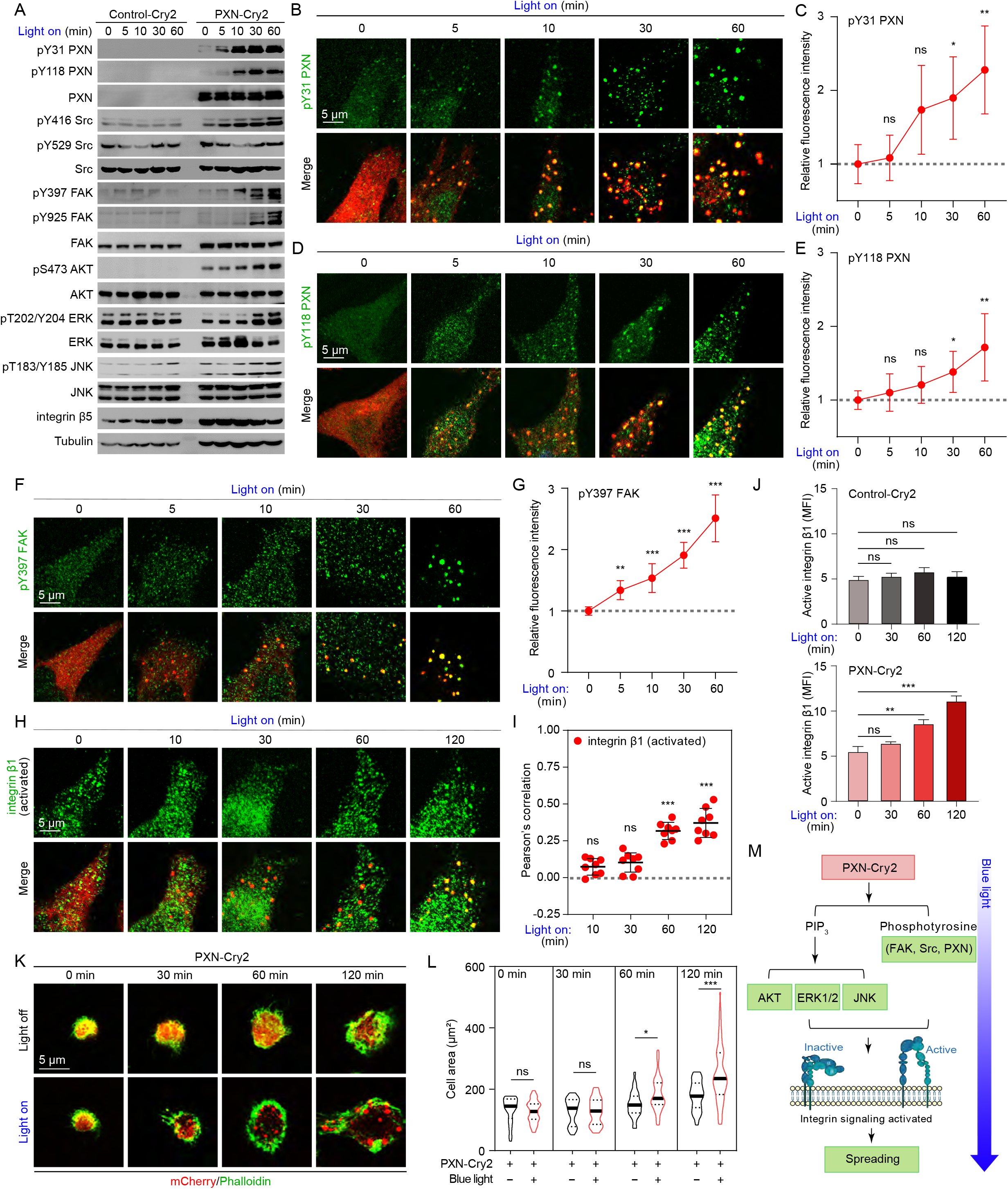
Light-induced PXN LLPS activates integrin signaling and accelerates cell spreading. (**A**) Lysates extracted from HeLa cells expressing either Control-Cry2 or PXN-Cry2 treated with blue light at different time points were separated on SDS-PAGE gels, and analyzed by immunoblotting with the indicated antibodies. The experiments were repeated 3 independent time. (**B**-**I**) HeLa cells expressing PXN-Cry2 were stimulated with blue light for different time periods. The cells were stained with the indicated antibodies in (**B**), (**D**), (**F**), (**H**). The relative fluorescence intensities of individual staining were quantified in (**C**), (**E**), (**G**), (**I**). Data are presented as mean ± SD. ns, not significant; *p < 0.05; **p < 0.01; ***p < 0.001 by one-way ANOVA. (**J**) *PXN*^-/-^ HeLa cells expressing Control-Cry2 or PXN-Cry2 were stimulated with blue light for different time periods and stained with antibody specifically against active integrin β1. The mean fluorescence intensity (MFI) was determined by flow cytometry. Data are shown as mean ± SEM, n = 3 independent experiments. ns, not significant; **p < 0.01; ***p < 0.001 by one-way ANOVA. (**K**) Representative confocal images of HeLa cells expressing PXN-Cry2 that were digested and plated on fibronectin-coated (5 μg/mL) cover slips in the absence (light off) or presence (light on) of blue light illumination for the indicated time. The cells were stained with Phalloidin to highlight morphology. (**L**) Quantification of the cell area from (**K**). ns, not significant; *p < 0.05; ***p < 0.001 by unpaired Student t-test. (**M**) Schematic summary of the results from (**A**-**L**).

### PXN LLPS facilitates FA formation

The feasibility to reconstitute functional FAs via tuning PXN LLPS implicates PXN as a core of FAs. To further assess the centrality of PXN in the protein network of FAs, we chose to contrast the biophysical and functional properties of a number of well-known FA-associated proteins. Among these, PXN and Zyxin represent LIM (Lin11, Isl-1, and Mec-3) domain containing proteins; FAK, Kindlin-2, and Talin-1 belong to proteins harboring the FERM (band 4.1, ezrin, radixin, moesin) domain; Src kinase contain typical SH domains, and several other proteins such as Vinculin, α-actinin, GIT1, and VASP, have their unique domain configurations (Fig. S3 A). We successfully obtained 9 of the 10 recombinant proteins in full length except Talin-1, due to its unusual large size (Fig. S3 B). Therefore, we only purified the FERM domain of Talin-1 instead (referred to as Talin-FERM). Largely consistent with prior reports (Case et al., 2022; Li et al., 2020; Zhu et al., 2020), several of these 10 proteins, including FAK, PXN, Zyxin, and GIT1 formed droplets in the physiological buffer conditions (150 mM NaCl, pH 7.5) in the absence of any crowding reagents (Fig. 4A). FAK exhibited the lowest threshold concentration (< 1 μM) necessary for LLPS, followed by PXN, GIT 1, and Zyxin (Fig. 4, A and B). Distinct from previous findings that PXN failed to condensate at up to 30 μM concentration(Case et al., 2022; Zhu et al., 2020), we observed that recombinant PXN forms condensates at ~ 5 μM concentration. This seemingly discrepancy is likely due to the fact that we purified PXN in the presence of 50 μM ZnSO_4_. And we found that zinc ions significantly potentiated PXN LLPS, which was mediated through the tandem zinc fingers locating within the C-terminus of PXN (data not shown). The rest of the FA-associated proteins failed to phase separate under the same conditions (Fig. S3 C). Next, we sought to explore if correlation exists between the LLPS potential and FA formation. We focused on FAK, PXN, Zyxin and GIT1 because these 4 proteins are most readily to phase separate under physiological buffer conditions. Upon interfering with the expression of individual protein in HeLa cells by siRNA (Fig. S3 D), we observed that FA formation was most susceptible to loss of PXN, followed by FAK (Fig. 4, C-E). Silencing Zyxin and GIT1 had no appreciable effects on the adhesion formation (Fig. 4, C-E). Together, these data suggest that LLPS propensity of FA proteins largely correlates with their roles in promoting FA assembly.

**Figure 4.**
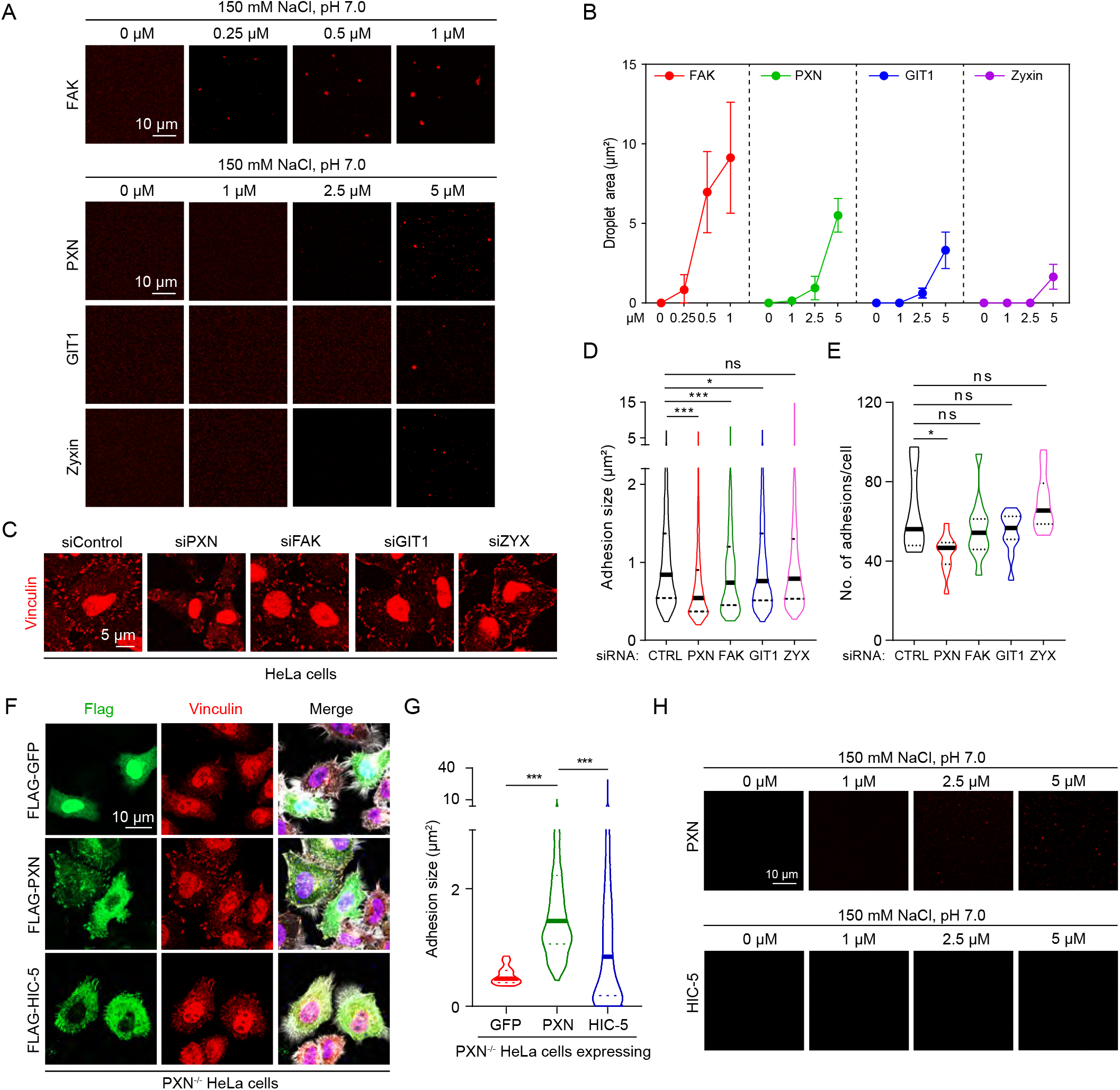
PXN undergoes LLPS and is required for FA formation. (**A**) LLPS of PXN, FAK, Zyxin, and GIT1 at the indicated concentrations in physiological buffer (150 mM NaCl, pH 7.5) in the absence of crowding agents. All the proteins were labeled with Cy3 NHS ester. Droplet size of each individual proteins formed at various concentrations was quantified in (**B**). (**C**) HeLa cells transfected with the indicated siRNA were fixed and stained with antibody against Vinculin. Representative images were shown from 2 independent experiments. Adhesion size and number were quantified and shown in (**D**) and (**E**), respectively. ns, not significant; *p < 0.05; ***p < 0.001 by one-way ANOVA. (**F**) *PXN*^-/-^ HeLa cells were transiently transfected with Flag-GFP, Flag-PXN, or Flag-HIC-5 constructs. The cells were stained with Phalloidin and antibody against Vinculin. The FA size under each experimental condition was quantified in (**G**). ***p < 0.001 by one-way ANOVA. (**H**) Representative images of LLPS of PXN and HIC-5 at the indicated protein concentrations.

Complete *PXN* deletion in HeLa cells almost ablated FA formation, which was efficiently rescued by reintroducing WT PXN. Nevertheless, we found that expression of HIC-5, the close paralogue of PXN, in the *PXN*^-/-^ cells could only marginally restore the defective FA formation (Fig. 4, F and G), This observation inspired us to compared the phase separation potentials of PXN with HIC-5. Indeed, whereas PXN reliably formed micro-sized droplets under the conditions we tested, HIC-5 failed to do so (Fig. 4 H and S4 E). These data also support our observation that Cry2-HIC-5 was unable to form blue light-induced condensates (Fig. 1, G and H). It is well-documented that while PXN family members share many of the same binding partners, a number of interactions appear to be specific to individual protein (Alpha et al., 2020). The distinct condensation potential and binding repertoire of PXN and HIC might explain why HIC-5 is not sufficient to rescue FA formation in the *PXN*^-/-^ HeLa cells.

To understand the structural basis of PXN LLPS, we generated N-terminal and C-terminal truncated proteins. The N-terminus was predicted to be mostly disordered (Fig. 5 A). *In vitro* LLPS assays showed that while the N-terminal IDR was sufficient to drive LLPS, albeit to a lesser extent compared to the full length PXN, C-terminal fragment formed amorphous aggregates under the same buffer conditions (Fig. 5 B and Fig. S4 A). PXN LLPS was potentiated by increasing ion strength (Fig. S4, B and C), indicating that electrostatic interactions are not the driving force of PXN LLPS, which is in line with the relatively uniform charge distribution alone the PXN primary sequence (Fig. 5 A). In contrast, 5% 1,6-hexanediol, which disrupts weak hydrophobic interactions, significantly impeded the PXN droplet formation (Fig. 5 C). The PXN droplets exhibited liquid-like attributes, including rapid fusion and fluorescence recovery following photobleaching (Fig. S4, D-F). Similarly, GFP-PXN ectopically expressed in the HeLa cells also forms discrete condensate structures, which were susceptible to 5% 1,6-hexanediol treatment (Fig. 5, D and E) and display liquid-like behavior, such as material exchange with the surrounding cytoplasm and continuous fusion (Fig. S4, G-I).

**Figure 5.**
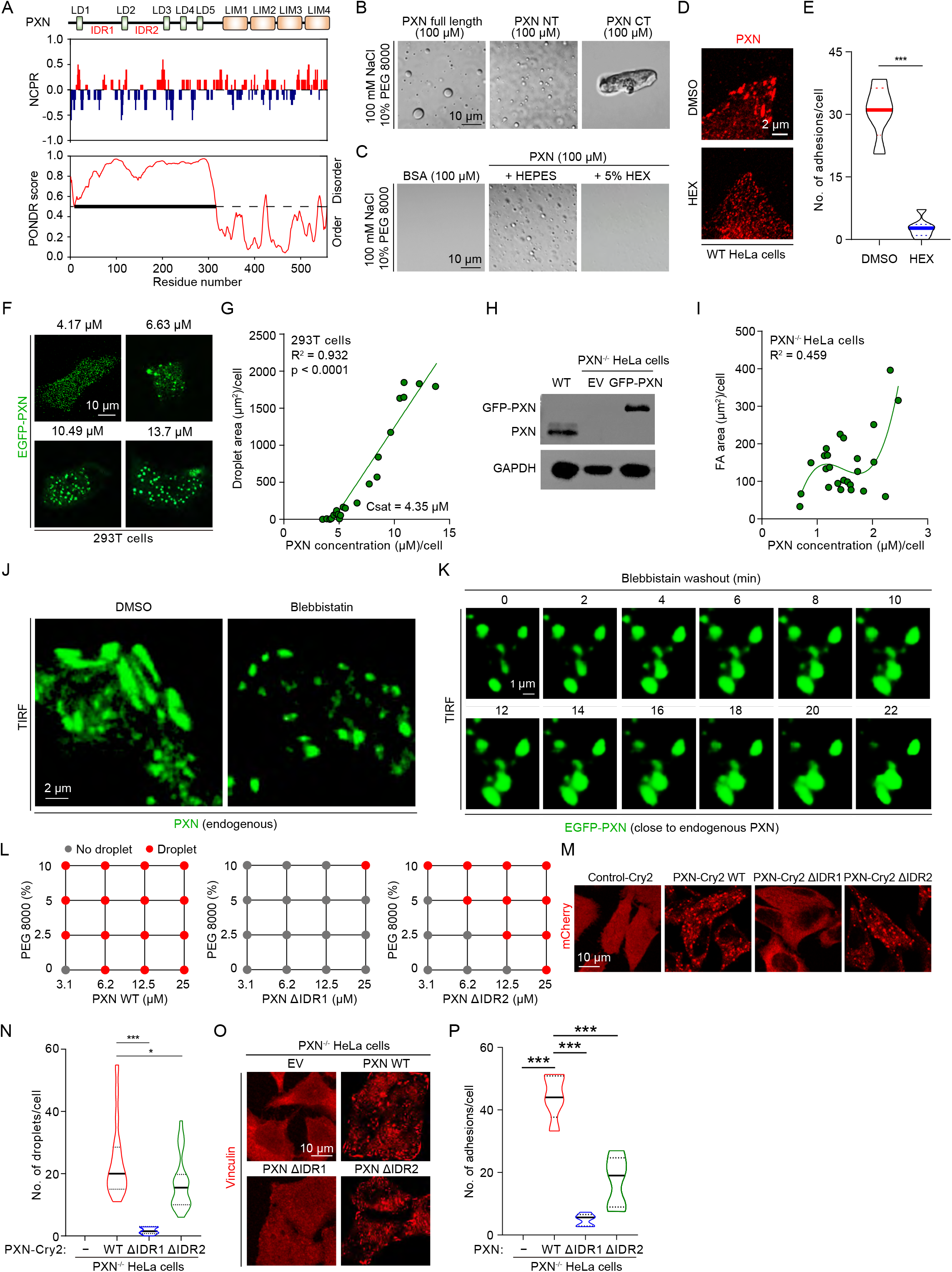
PXN LLPS promotes FA assembly. (**A**) Prediction of disordered regions (PONDR) and charge distribution (NCPR) were aligned with PXN domains. (**B**) Representative DIC images of LLPS of PXN full length, N-terminus (NT), and C-terminus (CT) *in vitro* (150 mM NaCl, pH 7.5, 10% PEG 8000). (**C**) Representative DIC images of PXN in the presence or absence of 5% 1,6-hexanediol. (**D**) WT HeLa cells were treated with 5% 1,6-hexanediol and stained with antibody against PXN. The number of PXN^+^ adhesions were quantified in (**E**). ***p < 0.001 by unpaired Student t-test. (**F**) Representative confocal images of EGFP-PXN expression at different concentrations in the 293T cells. The correlation between the protein concentration of EGFP-PXN and condensate area was shown in (**G**). (**H**) Protein samples prepared from WT HeLa and *PXN*^-/-^ HeLa cells reconstituted with empty vector (EV) or GFP-PXN were analyzed by immunoblotting against the indicated antibodies. (**I**) EGFP-PXN was reconstituted into *PXN*^-/-^ HeLa cells to near endogenous PXN level. The correlation between PXN concentration and FA area was determined. The graph was modeled with polynomial regression. (**J**) WT HeLa cells treated with DMSO or blebbistatin were stained with antibody against PXN and imaged with TIRF microscopy. (**K**) HeLa cells expressing EGFP-PXN close to the endogenous PXN expression were recovered from blebbistatin treatment and subjected to live cell imaging with TIRF microscopy. (**L**) Phase diagrams of PXN WT, ΔIDR1, and ΔIDR2 with increasing concentrations of PEG 8000. (**M**) Representative images of *PXN*^-/-^ HeLa cells expressing PXN-Cry2 WT, ΔIDR1, or ΔIDR2 stimulated with blue light for 10 min. Number of Opto-PXN droplets was quantified in (**N**). *p < 0.05; ***p < 0.001 by one-way ANOVA. (**O**) Representative images of *PXN*^-/-^ HeLa cells reconstituted with empty vector (EV), PXN WT, ΔIDR1, or ΔIDR2 immunostained with antibody against Vinculin. FA size was quantified and presented in (**P**). ***p < 0.001 by one-way ANOVA.

To delineate the condensation properties of PXN *in vivo*, we sought to determine the concentration above which PXN starts to demix. Towards this end, we chose to express GFP-PXN in the HEK293T cells because of the poor ability of these cells to form FAs (data not shown). We observed a protein concentration-dependent condensation of PXN (Fig. 5 F), and established a positive correlation between total PXN protein concentration and area of droplets (Fig. 5 G). The saturation concentration of PXN was determined to be 4.35 μM by quantitative fluorescent microscopy (Fig. 5 G), which highly agreed with the low micro-molar threshold concentration (~ 5 μM) of PXN phase separation *in vitro* (Fig. 4 A). Taken together, the overexpression experiments indicate that PXN harbors an intrinsic capacity to form phase separated compartments. To assess the condensation property of PXN at the endogenous level, we reconstituted GFP-PXN into *PXN*^-/-^ HeLa cells and carefully sorted the cells such that GFP-PXN expression was close to endogenous PXN levels (Fig. 5 H). Under this condition, PXN formed condensates at less than 1 μM protein concentration, which was much lower than the threshold of spontaneous condensation in the 293T cells (Fig. 5 I). The protein concentration of PXN was not linearly correlated with the area of FAs (Fig. 5 I), which implicates active roles of additional factors in governing PXN condensation under physiological conditions. Together, these data suggest that PXN favors phase separation initiated by homotypic interactions at exogenous levels, but undergoes complex condensation at the endogenous levels. If this hypothesis is true, the client proteins of FAs should lower the threshold concentration of PXN to phase separate. Indeed, several of these client proteins including Src, Vinculin, Kinlin-2 and GIT-1 significantly promoted PXN phase separation (Fig. S4 J). Our findings are consistent insofar as recent report showing that FAK and p130Cas synergistically phase separate to promote phase separation, integrin clustering and FA formation in the MEFs (Case et al., 2022).

When comparing the fusion dynamics of phase-separated PXN droplets *in vitro* (Fig. S4 D) and PXN at the FAs (Fig. S4 G), we noticed that the PXN fusion *in vivo* were dramatically slower and more constrained than that of a liquid phase. This observation implicates additional interactions not present in the condensed PXN droplets, such as protein-lipid interactions and actomyosin contraction, take place at the mature FAs. To directly test the role of actomyosin contraction in modulating the material properties of FAs, we treated cells with blebbistatin, which reversibly inhibits Myosin II ATPase activities. As expected, endogenous PXN-enriched adhesions exhibited morphological changes from rod-like to spherical droplet-like after blebbistatin treatment (Fig. 5 J). Upon blebbistatin withdrawal, the EGFP-PXN^+^ adhesions gradually elongated and fused into large sub-apical membrane compartments as revealed by live cell imaging by TIRF microscopy (Fig. 5 K). These data suggest that endogenous PXN compartments behave as a viscoelastic fluid, which grow on the sub-apical membrane, and are modulated by F-actin, among other factors.

To dissect the role of PXN phase separation in FA formation, we deleted the largest disordered regions locating between LD1 and LD2, and LD2 and LD3 (referred to as ΔIDR1, and ΔIDR2, respectively). Interestingly, deleting IDR1 almost completely abolished the ability of PXN to form droplets in the *in vitro* phase separation assay (Fig. 5 L and Fig. S4 K). Consistently, PXN-Cry2 ΔIDR1 failed to undergo blue-light dependent phase separation in the cells (Fig. 5, M and N). *PXN* deletion in HeLa cells significantly impeded FA formation, which was efficiently rescued by reintroducing WT PXN. Expression of PXN ΔIDR1 mutant was incapable of reversing the FA assembly defects of PXN null cells (Fig. 5, O and P). In contrast, PXN ΔIDR2 mutant retained ability to phase separate, form light-induced condensates, and promote FA formation, although to a lesser degree compared to the WT PXN (Fig. 5, L-P). The markedly distinct functions of PXN WT, ΔIDR1, and ΔIDR2 in FA formation were unlikely due to changes in binding to its interacting proteins because these PXN variants were capable of interacting with FAK, Src, Vinculin, and Kindlin-2, almost equally (Fig. S 4L). Taken together, these data suggest that PXN promote FA assembly/maturation through phase separation.

### Specific and non-specific molecular interactions cooperatively regulate FA assembly and integrin signaling

Although our data implicates the important role of the complex condensation of PXN in assembling biological adhesions, the nature of forces that drives this process remains unknown. Prior studies have isolated LD2 and LD4 domains to be the key protein-protein interacting motifs of PXN. For example, PXN was reported to interact with FAK through LD2 and LD4 (Brown et al., 1996; Turner et al., 1999). It also interacts with Vinculin, Talin-1, and GIT1 via the LD2 and/or LD4 domains (Brown et al., 1996; Turner et al., 1999; Zacharchenko et al., 2016). Therefore, we generated a PXN mutant lacking the LD2 and LD4 motifs, which is referred to as ΔLD2/4 and assessed its impact on PXN phase separation properties and functions. As the first step, we confirmed that LD2/4 deletion abolished the specific molecular interaction between PXN and its well-known binding proteins, including FAK, Vinculin, and GIT1 (Fig. 6 A). Expectedly, Kindlin-2, which was reported to interact with the LIM3 domain of PXN (Theodosiou et al., 2016), still remained bound to the ΔLD2/4 mutant (Fig. 6 A). Interestingly, we found that PXN also hetero-oligomerized with HIC-5 through LD2 and/or LD4 (Fig. 6 A). Next, we purified the full length PXN as well as the ΔLD2/4 mutant, and performed *in vitro* LLPS assays (Fig. S5 A). We observed that PXN without LD2 and LD4 domains still retained the ability to phase separate, although requiring higher threshold concentration (Fig. 6 B). Next, we constructed PXN-Cry2 ΔLD2/4, and reconstituted it into *PXN*^-/-^ HeLa cells (Fig. S5 B). Using PXN-Cry2 WT as a control, we found that the Cry2-mCherry-PXN ΔLD2/4 mutant was capable of forming light-induced Opto-PXN droplets, albeit to a lesser degree compared to PXN-Cry2 WT (Fig. S5, C and D). Intriguingly, the PXN interacting proteins, including FAK, Vinculin, Talin-1, and GIT1 still exhibited selective enrichment in the Opto-PXN droplets formed by ΔLD2/4 mutant (Fig. 6, C and D). More importantly, phosphorylation of FAK at Y397, and PXN at Y31 and Y118 was increased in cells expressing the ΔLD2/4 mutant PXN upon 60 min of blue light stimulation (Fig. 6, E-G; and Fig. S5 E). Note that the increase in the phosphorylation of FAK and PXN in cells expressing ΔLD2/4 mutant PXN did not reach the same degree compared to those expressing WT PXN. We also reconstituted the WT and ΔLD2/4 mutant PXN into *PXN*^-/-^ HeLa to examine the potential functional consequences. As was observed in the Opto-PXN system, the ΔLD2/4 mutant prominently colocalized with the FA-associated proteins including HIC-5, Vinculin, etc., and promoted FA formation, but not as efficiently as the WT PXN (Fig. 6, H and I; and Fig. S5 F). These data imply that complex condensation involving both specific and non-specific intermolecular interactions between PXN and client proteins facilitate FA assembly/maturation. PXN depletion in the HeLa cells profoundly accelerated cell migration as indicated by the widely used transwell assays, which was significantly reversed by exogenously expressed WT. ΔLD2/4 PXN inhibited cell migration as potently as that of WT PXN (Fig. S5, G and H), which was entirely consistent with the largely normal integrin signaling in cells expressing ΔLD2/4 PXN (Fig. 6 A). Taken together, our data highlight an unexpected yet essential role of non-specific molecular interactions, that underlies complexes condensation of PXN, in regulating adhesion assembly and integrin signaling.

**Figure 6.**
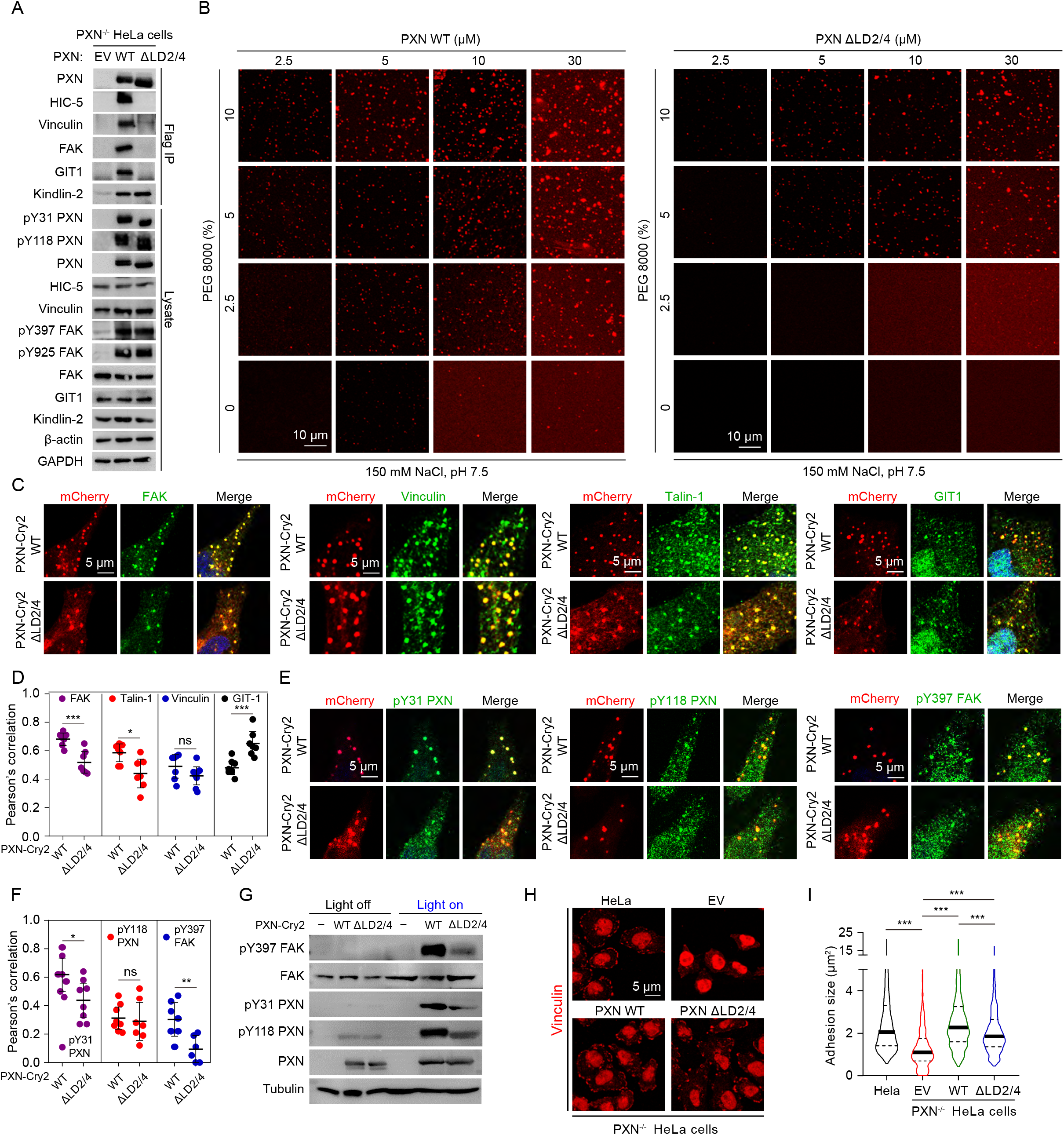
Specific and non-specific molecular interactions cooperatively regulate FA assembly and integrin signaling. (**A**) *PXN*^-/-^ HeLa cells reconstituted with empty vector (EV), Flag-tagged PXN WT, or ΔLD2/4 were subjected to immunoprecipitation with antibody against Flag. The protein samples were then analyzed by immunoblotting against the indicated antibodies. (**B**) Representative images of LLPS of PXN WT and ΔLD2/4 with increasing concentrations of PEG 8000. (**C**-**F**) HeLa cells expressing PXN-Cry2 WT or ΔLD2/4 were activated with blue light, fixed and immunostained with the indicated antibodies. Representative confocal images were shown in (**C**), and (**E**). Quantification of Pearson’s correlation of each individual protein with the Opto-PXN droplets formed by PXN-Cry2 WT or ΔLD2/4 were shown in (**D**), and (**F**). Data are shown as mean ± SD. ns, not significant; *p < 0.05; **p < 0.01; ***p < 0.001 by unpaired Student t-test. (**G**) HeLa cells expressing with PXN-Cry2 WT or ΔLD2/4 were treated with blue light for 60 min and harvested. Lysates were analyzed by immunoblotting with the indicated antibodies. (**H**) Representative confocal images of *PXN*^-/-^ HeLa cells reconstituted with empty vector (EV), Flag-tagged PXN WT, or ΔLD2/4 immunostained with antibody against Vinculin. FA size was quantified and presented in (**I**). ***p < 0.001 by one-way ANOVA.

## Discussion

To our knowledge, our light-inducible synthetic approach is the first to achieve designer adhesion complexes that are compositionally and functionally similar to those in human cells. Harnessing a single disordered fragment of the key scaffold protein of FAs, PXN, we developed a simple, elegant tool to efficiently reconstitute FA-like structures *in cellulo*. Although our Opto-PXN system was inspired by the OptoDroplets originally invented by Shin *et al* (Shin et al., 2017), Opto-PXN is substantially different from OptoDroplets in that Opto-PXN is sufficient to reconstitute complex biological structures where as OptoDroplets do not. This distinction makes sense when considering PXN is centrally located in the network of FAs, but proteins like FUS, which are harnessed in the OptoDroplets, are client proteins of multiple biomolecular condensates. Therefore, our Opto-PXN provides a unique platform to delineate the mechanisms underlying adhesion formation, growth and maturation.

Build on our observation and several decades of mechanistic studies on the dynamics of FAs, we propose the following simplified 3-step model to depict FA assembly. First, PXN are recruited to nascent adhesions. Selective enrichment of PXN within this sub-apical membrane domain might be sufficient to trigger PXN LLPS. Subsequently, the phase separated PXN compartments specifically sequester FA-associated proteins. In the physiological settings, specific and non-specific intermolecular interactions function cooperatively to facilitate the recruitment of FA-associated proteins by PXN. Finally, protein-lipid interaction, actin cytoskeleton and other yet unknown mechanisms function cooperatively to induce molecular reorganization such that the liquid-like nascent adhesion mature into viscoelastic fluid-like mature adhesions. Importantly, we must emphasize that we have focused on the roles of PXN LLPS, which is intuitively consistent with many features of the cytoplasmic macromolecular complex of FAs, our data nevertheless don’t exclude the well-established roles of other factors such as extracellular ligand binding, protein-lipid interaction, and the actomyosin cytoskeleton in FA assembly/maturation. Instead, we propose that protein-driven LLPS cooperates with these aforementioned factors to regulate different aspects of integrin signaling transduction, nascent adhesion formation and adhesion maturation.

Several FA-resident proteins have been demonstrated to harbor LLPS property (Case et al., 2022; Li et al., 2020; Wang et al., 2021; Zhu et al., 2020). In fact, our own experimental data also show that many, if not all, of the adhesion proteins are prone to phase separate under the “right” conditions (i.e. high protein concentration, crowder, etc.). This observation however, raises an exceptionally important question: what’s the exact relationship between LLPS and FA dynamics? Our data place PXN at the central node of the protein network of FAs and prove that PXN LLPS facilitates FA assembly. Although we focused on PXN, other proteins with phase separation traits might also contribute to FA assembly/maturation. In fact, LIMD1, which is structurally similar to that of PXN, was recently shown to undergo phase separation and force-dependent localization to adhesions during maturation (Wang et al., 2021). Future work will be needed to understand how protein LLPS might alter the compositional and biophysical properties of FAs and whether additional proteins, similar to PXN and LIMD1, become more important at different stages of FA formation and maturation.

It is noteworthy that unlike HeLa cells, FA formation is largely normal in the PXN null MEFs (Hagel et al., 2002). We speculate the existence of multiple “scaffolds” in the MEFs. Therefore, the “scaffold” role of PXN to initiate compartmentation of adhesions is likely context-dependent and can be compensated by other “scaffolds” under certain conditions. In fact, evidence has just emerged that FAK and p130CAS also synergistically promotes integrin clustering and FA formation via phase separation in the MEFs (Case et al., 2022). Likewise, multiple “scaffolds” seem to work in parallel in other membrane-less systems, such as stress granules (Cirillo et al., 2020; Yang et al., 2020). Understanding the interplay between different scaffolds will shed in-depth insights into the organizing principles of the membrane-less organelles.

## Materials and methods

### Plasmids

Plasmids were constructed using the 2 × MultiF Seamless Assembly Kit (Abclonal; RK21020). Vector backbones were linearized by DNA restriction endonucleases. The target fragments were amplified by high-fidelity DNA polymerase 2 × Phanta Max Master Mix (Vazyme; P515-01). The PCR primers were designed using the SnapGene software, including 20 bases overlapping with the vectors at the 5’end. The cloning was performed according to the manufacturer instructions. Sanger sequencing was performed to confirm the correct clones.

To generate retroviral constructs, cDNA was amplified using a standard PCR-based approach. The amplified cDNA was cloned into pLV retroviral vector using 2 × MultiF Seamless Assembly Kit. The Flag-*PXN* plasmids were constructed by insert into the pCDNA3.3 vector by Seamless Assembly Kit. Constructs containing the *PXN* IDR (*IDR-PXN-mCherry-Cry2*) were generated by inserting the *PXN* IDR fragment into the Lenti-*mCherry-Cry2* backbone (a generous gift from Dr. Shuguo Sun, Huazhong University of Science and Technology) by Seamless Assembly Kit. Constructs carrying point mutations were generated using overlap PCR. The mutant fragments were amplified by high-fidelity DNA polymerase 2 × Phanta Max Master Mix followed by DpnI (New England BioLabs; R0176S) digestion in 37°C for 1 hour to eliminate the templates, and the PCR products were used to transform DH5α competent cells (Shenzhen KangTi Life Technology; KTSM101L). Sanger sequencing was performed to confirm the correct clones.

### Generation of knock-out cell lines with CRISPR/Cas9

*PXN* knockout cells were created through the CRISPR/Cas9 technology. The guide RNA sequences were designed using online tool the Optimized CRISPR Design (https://portals.broadinstitute.org/gppx/crispick/public). The guide sequence was 5’-ATCCCGGAACTTCTTCGAGC-3’ for human *PXN* and for 5’-TTGAGGGCCTCGCTGGACGG-3’ mouse *Pxn*. Cells were transiently transfected with PX459 (Addgene; #48139) and selected with puromycin for 2 days. The pool was scattered into a 10 cm petri dish. Single clones were picked up and expanded for immunoblotting analyses for the absence of PXN.

### Cell culture, transient transfection and lentivirus infection

The 293T (CRL-1573), and HeLa (CCL-2) cell lines were purchased from (American Type Culture Collection; ATCC), tested negative for mycoplasma contamination. All cells were grown in DMEM(L120KJ), supplemented with 10% FBS (Moybio; S450), penicillin/streptomycin (BasalMedia; S110JV), and Glutamax (BasalMedia; S210JV) at 37°C (5% CO_2_). To transiently overexpress cDNA constructs, cells were transfected with Polyethyleneimine Linear (BIOHUB; 78PEI25000) or Lipofectamine 2000 (Thermo Fisher; 11668019) according to the manufacturer’s instructions. Knockdown experiments were performed with Lipofectamine RNAi Max (Life Technologies; 13778075) according to the manufacturer’s instructions. The following siRNAs synthesized by GenePharma (Shanghai, China) were used: si*PXN*: 5’-CCCUGACGAAAGAGAAGCCUAUU-3’ and 5’-UAGGCUUCUCUUUCGUCAGGGUU-3’; si*FAK*: 5’-GCGAUUAUAUGUUAGAGAUAGUU-3’ and 5’-CUAUCUCUAACAUAUAAUCGCUU-3’; si*GIT1*: 5’-CCUUGAUCAUCGACAUUCUTT-3’ and 5’-AGAAUGUCGAUGAUCAAGGTT-3’; si*ZYX*: 5’-GCCUCAGGUCCAACUCCAUTT-3’ and 5’-AUGGAGUUGGACCUGAGGCTT-3’. To generate HeLa cells stably expressing wild-type or mutant forms of PXN, lentivirus was produced by co-transfecting 293T cells with psPAX2 (Addgene; #12260), pMD2.G (Addgene; #12259) and pLV retroviral vectors containing different *PXN* cDNAs at 70%–80% confluency. Supernatants were harvested twice at 48 and 60 hours after transfection, and centrifuged at 8000 r.p.m. for 3 min at room temperature (RT) to remove cell debris followed by passage through a 0.22 μm filter. The indicated cells were incubated with retroviral supernatant containing 1:1000 dilute of polybrene (Santa Cruz Technology; sc-134220) for 1 day, and the transduced cells were FACS-sorted by the presence of GFP or selected with antibiotics.

### Immunoblotting

Cells were harvested and lysed with ice cold lysis buffer (20 mM Tris-HCl, pH 7.5, 150 mM NaCl, 1 mM EDTA, 1 mM EGTA, 1% Triton X-100, 2.5 mM sodium pyrophosphate, 1 mM ß-glycerolphosphate, protease inhibitor cocktail). The lysates were then centrifuged to clear cell debris. The supernatant was then prepared with an equal amount of 2 × SDS sample buffer, and electrophoretically separated on SDS-PAGE gels. Proteins were transferred to PVDF membranes. After blocking with 5% skin milk, the membranes were probed with the following primary antibodies: rabbit anti-Zyxin (GeneTex; GTX132295), rabbit anti-phospho-PXN (Y31) (Invitrogen; 2024882), rabbit anti-phospho-PXN (Y118) (Cell Signaling Technology; 2541S), rabbit anti-PXN (Proteintech; 10029-1-Ig), rabbit anti-PXN (Abcam; ab32084), rabbit anti-Src (Cell Signaling Technology; 2109S), rabbit anti-phospho-Src (Y416) (Cell Signaling Technology; 59548S), rabbit anti-phospho-Src (Y527) (Cell Signaling Technology; 2105T), rabbit anti-FAK (Cell Signaling Technology; 3285S), rabbit anti-phospho-FAK (Y397) (Abcam; ab81298), rabbit anti-phospho-FAK (Y925) (Abcam; ab38512), rabbit anti-Pan-AKT (Abclonal; A18675), rabbit anti-phospho-AKT (S473) (Cell Signaling Technology; 4060), rabbit anti-ERK1/2 (Abclonal; A4782), rabbit anti-phospho-ERK (T202/Y204) (Cell Signaling; 4370), rabbit anti-JNK1/2/3 (Abclonal; A4867), rabbit anti-phospho-JNK (T183/Y185) (Cell Signaling Technology; 4668), rabbit anti-HIC-5 (Proteintech; 10565-1-AP), mouse anti-GIT1 (BD Transduction; 611396), rabbit anti-Kindlin-2 (Proteintech; 11453-1-AP), mouse anti-α1 Na^+^/K^+^ ATPase (Abcam; ab7671), rabbit anti-α-tubulin (Cell Signaling Technology; 2125S), mouse anti-GAPDH (Santa Cruz Biotechnology; sc-32233), mouse anti-actin (Proteintech; 66009-1-Ig), rabbit anti-integrin β5 (Cell Signaling Technology; 3629S). The membranes were then incubated with HRP-conjugated secondary antibodies (Jackson ImmunoResearch; 115-035-003, 111-035-003), and bands were detected using chemiluminescence detection kit (Merck Millipore; WBKLS0050).

### Immunoprecipitation

Total cell lysates were prepared from cells using ice cold lysis buffer, and anti-DYKDDDDK G1 Affinity Resin (GenScript; L00432-10) were then added into the cleared lysates for 3 hours at 4°C. The beads were spun and washed with lysis buffer 3 times. The beads were boiled for 10 min in 2 × SDS sample buffer followed by standard immunoblotting procedure as described above.

### Immunofluorescence

Cells grown on glass coverslips were cultured to 60-80% confluence. Cells were fixed with 4% formaldehyde (diluted in PBS) at RT for 10 min, permeabilized with 0.1% Triton X-100 (diluted in PBS), blocked with 3% BSA, and incubated with primary antibodies overnight at 4°C. The following primary antibodies were used: mouse anti-Flag (GNI; GNI4110-FG), rabbit anti-Vinculin (Proteintech; 26520-1-AP), mouse anti-G3BP1 (BD Transduction; 611126), rabbit anti-YBX1 (Abclonal; A7704), rabbit anti-HIC-5 (Proteintech; 10565-1-AP), mouse anti-HIC-5 (BD Transduction; 611164), rabbit anti-FAK (Cell Signaling Technology; 3285S), rabbit anti-Talin-1 (Proteintech; 14168-1-AP), mouse anti-GIT1 (BD Transduction; 611396), rabbit anti-Kindlin-2 (Proteintech; 11453-1-AP), rabbit anti-LIMD1 (Proteintech; 28106-1-AP), rabbit anti-α-actinin (Proteintech; 11313-2-AP), rabbit anti-VASP (Proteintech; 13472-1-AP), mouse anti-TNS3 (Santa Cruz Biotechnology; sc-376367), rabbit anti-Zyxin (GeneTex; GTX132295), mouse anti-integrin α2β1 (Abcam; ab24697), rat anti-integrin β1 mAb13 (Abcam, MABT409), mouse anti-integrin β1 (12G10) (Abcam; ab30394), rabbit anti-PXN (Proteintech; 10029-1-Ig), and rabbit anti-PXN (Abcam; ab32084). The cells were then incubated with Alexa-Fluor-conjugated secondary antibody (Jackson ImmunoResearch; 115-545-003, 115-585-003) for 1 hour at RT in the dark, and mounted on slides with mounting media (SouthernBiotech; 0100-01). Cells were imaged by Zeiss LSM 900 confocal microscopy with a 63x oil objective. All the confocal images were obtained by focusing on the middle of the cells unless otherwise specified.

### Blue light treatment

Cells were illuminated using Leica M165 FC fluorescent stereo microscope in a humidified chamber with 5% CO_2_ for continuous blue light stimulation.

### Protein expression and purification

The cDNAs of FA proteins were cloned into a modified version of a PET-16b vector. The template vector was engineered to include a 6 x His-MBP-TEV followed by either GFP or mCherry using 2 x MultiF Seamless Assembly Method according to the manufacture’s instructions. All expression constructs were sequenced to ensure sequence accuracy. For protein expression, plasmids were transformed into BL21(DE3) *E. coli* cells (AngYu; G6030-10). A single colony was inoculated into LB media containing ampicillin and grown in LB media to an optical density of 0.6-0.8 at 37°C, followed by overnight induction with 1 mM isopropyl-b-D-thio-galactopyranoside (IPTG) at 16°C. Cells were pelleted and resuspended in binding buffer (20 mM Tris pH 7.5, 500 mM NaCl) and lysed using a homogenizer. The lysates were cleared by centrifugation and purified by Ni-NTA agarose (Nuptec; NRPB57L-100). The proteins were eluted with elution buffer (20 mM Tris pH 7.5, 500 mM NaCl, 300 mM imidazole). Eluate was concentrated and cleaved with TEV protease at 4°C. The cleaved tags and TEV protease were separated from the protein samples by a second round of Ni-NTA affinity chromatography followed by a Sephadex 200 size-exclusion column in 20 mM Hepes, 100 mM NaCl, pH 8.0. The collected fractions were then verified with SDS–PAGE. Purified fractions were pooled, concentrated, and flash-frozen in liquid nitrogen.

The cDNAs of human full length FAK was subcloned into the pMLink vector with a N-terminal 8 × His tag. FAK protein expression was done by Expi293F cells (Invitrogen; A14527). The Expi293F cells were grown in Union-293 media (Union-Biotech, Shanghai) under 5% CO_2_ in a ZCZY-CS8 shaker (Zhichu Instrument) at 37°C with 120 r.p.m. When the cell density reached 2.0 × 10^6^ cells/mL, pMlink-FAK was transiently transfected into the cell with polyethyleneimines (PEIs) (Polysciences). The transfected cells were cultured at 37°C for 48 hours before harvesting. The harvested cells were suspended in lysis buffer containing 50 mM Tris-HCl at pH 8.0, 150 mM NaCl, 20 mM Imidazole and 10% glycerol. Cells were lysed by a high-pressure homogenizer and cleared by centrifugation. FAK was purified using Ni Sepharose High performance resin (Cytiva), followed by gel filtration using Superdex Increase 200 10/300 (Cytiva) with buffer containing 50 mM HEPES, 500 mM NaCl, 1 mM DTT. Purified proteins were concentrated and buffer-exchanged in Amicon filters (Millipores).

### Protein labelling

Fluro 488 NHS ester (AAT Bioquest; 1810) or Cy3 NHS ester (AAT Bioquest; 271) were dissolved in DMSO at a concentration of 10 mg/mL and incubated with the corresponding proteins at 1:1 (molar ratio) at RT for 1 hour with continuous stirring. Free dye was removed using a desalting column (Merck Millipore; UFC501024), the labeled proteins were aliquoted and flash-frozen in liquid nitrogen.

### *In vitro* LLPS assay

Droplet formation of purified protein was monitored by fluorescence and differential interference contrast (DIC) microscopy using a confocal microscope (Zeiss LSM 900). Mixtures in a total solution volume of 2 μL were placed in on slides with double-sided tape. Cover glasses were placed on top to seal the slides. Imaging was performed on a Zeiss LSM900 microscope, using a plan-apochromat × 63/1.4 oil DIC M27 objective.

### Fluorescence recovery after photo-bleaching (FRAP)

FRAP analyses were performed using a Zeiss LSM 900 laser scanning microscope equipped with a HC PL APO 100×/1.40 NA oil CS2. Intracellular assemblies were bleached in a 1-2 μm^2^ circular region of interest (ROI) using a 3.25 s pulse of the 405 nm laser line at full power. Recovery was monitored every 0.65 s for 400 frames. Droplets formed in vitro were bleached in a circular 0.2 μm^2^ region of interest using a 0.5 s pulse of the 488 nm laser line at full power. Recovery was monitored every 0.5 s for 180 frames. Recovery curves were analyzed using Metamorph. Plotting and curve fitting were carried out in GraphPad Prism 8.0.

### Quantification of protein concentration *in vivo*

To determine the saturation concentration of GFP-PXN condensation in the HEK293T or HeLa cells, we purified GFP protein *in vitro* and measured the protein concentration by BCA assay. We then photographed the fluorescence of GFP protein or GFP-PXN in the cells at different concentrations under the same parameters with Z-Stack, and then fitted the average fluorescence intensity of GFP protein and the corresponding concentrations to obtain a standard curve. We used Imaris to quantify the average fluorescence intensity of GFP-PXN in cells, and calculated the concentration of GFP-PXN in HeLa or HEK293T cells based on the standard curve. The droplet area in the HEK293T cells and FA area in the HeLa cells were quantified by Imaris, then were plotted against the total protein concentration of GFP-PXN.

### Flow cytometry

Adherent HeLa cells were fixed with 2% PFA, and resuspended in MACS buffer (PBS, 0.5% BSA, 2 mM EDTA, pH 7.2). The samples were incubated with Fc block for 15 min, followed by centrifuge to remove supernatant. After incubation with primary antibody for 1 hour, the samples were washed 3 times with MACS buffer before incubation with second antibody for 30 min. The cells were then washed 3 times with MACS buffer, resuspended in MACS buffer for flow cytometry (BD Fortessa X-20).

### Plasma membrane fractionation

The plasma membrane fractions were prepared using the ProteoPrep Membrane Extraction Kit (Sigma-Aldrich; PROT-MEM) according to the manufacturer’s instructions.

### Transwell assay

30,000 cells were seeded in the top chambers of the transwell plates (Corning Incorporated; 3422) in FBS-free media with membrane inserts without matrigel coating. DMEM supplemented with 10% FBS were added to the lower compartment of the plates as attractant. After incubation at 37°C for 16-18 hours, cells on the upper surface of the membranes were removed with cotton swabs, and the cells that migrated to the lower surface were fixed by 4% PFA (Leagene; DF0135) for 10 min and stained with 0.2% (w/v) crystal violet (Sangon Biotech; A600331-0100) for 10 min and washed with ddH_2_O. The stained cells were imaged by Olympus IX51 4 x objective. All migration assays were repeated at least 3 times.

### Quantification and statistical analyses

Quantification of droplet size from *in vitro* LLPS assays was done by manual image segmentation (Image J). Briefly, images were converted to 8-bit format. The converted images were then Gaussian filtered to reduce background noise and the threshold was adjusted based on the intensity. Detected objects in each image were examined manually to confirm their validity. Only droplets in the focal plane (identified as having a thin, low-contrast droplet border) were considered.

Pearson correlation coefficient of 2 proteins was determined by Coloc2 Plugin of Image J.

Measurements of FA area were done with the Imaris software (Bitplane) using the surface reconstruction tool.

All analyses were performed blinded to genotype. All quantitative data were shown as mean ± SEM from n ≥ 3 biological replicates unless otherwise specified. Statistical significance was determined by Student’s t tests or ANOVA as appropriate, and *p < 0.05 was considered statistically significant. Statistical parameters are also reported in the figures and legends.

## Acknowledgements

We are grateful to Dr. Shuguo Sun (Huazhong University of Science and Technology), and Peipei Zhang (Peking University) for providing plasmids. We thank Dr. Pilong Li (Tsinghua University), Drs. Mondira Kundu and J. Paul Taylor (St Jude Children’s Research Hospital), and Dr. Xuebiao Yao (University of Science and Technology of China) for valuable discussions. All the imaging data were acquired in the Core Facility of Biomedical Sciences, Xiamen University. This work was supported by the National Natural Science Foundation of China (32071235 to B.W.), Guangdong Basic and Applied Basic Research Foundation (2020A1515111186 to B.W.), and National Key R&D Program of China (2021YFC2100100 to Q.X.).

## Author contributions

B.W. and P.L. designed the experiments. P.L., Y.W., S.Z., J.Z., S.Y., J.W., S.M., M.Z., Z.G., Q.L., W.J., Q.X., and B.W. performed experiments and/or analyzed the data. B.W. wrote the manuscript. Q.X. edited the manuscript.

## Figure legends

**Figure S1**. Characterization of light-induced Opto-PXN condensates (related to Figure 1 and Figure 2). (**A**) Representative photographs of WT and *PXN*^-/-^ HeLa cells stained with antibody against Vinculin. (**B**) *PXN*^-/-^ HeLa cells stably expressing PXN-Cry2 were subjected to the indicated paradigm of blue light treatment. Representative images are shown from 3 independent experiments. (**C**) HeLa cells expressing PXN-Cry2 were pre-treated with 5% 1,6-hexanediol for 10 min and then stimulated with blue light for 10 min. (**D**) Lysates prepared from WT HeLa cells stably expressing empty vector (EV), PXN-Cry2, or HIC-5-Cry2 were analyzed by immunoblotting against the indicated antibodies. (**E**, **F**) Representative images of *PXN*^-/-^ HeLa cells expressing either PXN-Cry2 or Control-Cry2 immunostained with the indicated antibodies. (**G**-**J**) *PXN*^-/-^ HeLa cells expressing PXN-Cry2 were activated with blue light for the indicated time, and immunostained with antibodies against G3BP1 (**G**) and YBX1 (**I**), respectively. The Pearson’s correlation of G3BP1 and YBX1 with the Opto-PXN droplets were quantified and presented as mean ± SD in (**H**, **J**). (**K**-**N**) WT HeLa cells transiently transfected with Opto-FUS were stimulated with blue light for the indicated time. The cells were then fixed and immunostained with antibodies against PXN or HIC-5. Representative confocal images were shown in (**K**), and (**M**). And the Pearson’s correlation of PXN or HIC-5 with Opto-FUS condensates were quantified in (**L**), and (**N**). ns, not significant by one-way ANOVA. (**O**) WT HeLa cells expressing PXN-Cry2 were immunostained with the indicated antibodies before and after blue light stimulation. Representative confocal images were shown.

**Figure S2**. Light-induced PXN LLPS activates integrin signaling (related to Figure 3). (**A**) The relative band intensities of Figure 3A were quantified by densitometry. Data are presented as mean ± SEM, n = 3 independent experiments. (**B**-**C**) HeLa cells expressing PXN-Cry2 were illuminated with blue light for the indicated time, and immunostained with antibodies against inactive integrin β1. The Pearson’s correlation of inactive integrin β1 and Opto-PXN was quantified in (**C**). Data are presented as mean ± SD. ns, not significant by one-way ANOVA. (**D**) *Pxn*^-/-^ MEFs expressing PXN-Cry2 were activated with blue light for the indicated time, and immunostained with the indicated antibodies. (**E**) *Pxn*^-/-^ MEFs expressing Control-Cry2 or PXN-Cry2 were treated with blue light for 2 h. The cells were harvested and analyzed by immunoblotting using the indicated antibodies. (**F**) Representative confocal images of HeLa cells expressing Control-Cry2 that were digested and plated on fibronectin-coated (5 μg/mL) cover slips in the absence (light off) or presence (light on) of blue light for the indicated time. The cells were stained with Phalloidin to highlight morphology. (**G**) Quantification of the cell area from (**F**). ns, not significant by unpaired Student t-test.

**Figure S3**. PXN undergoes LLPS and is required for FA assembly (related to Figure 4). (**A**) Domain organization of representative adhesion proteins. (**B**) Recombinant FA proteins were separated by 10% and 12% SDS-PAGE gels, and visualized by Coomassie Blue. (**C**) LLPS of representative adhesion proteins at the indicated concentration in physiological buffer (150 mM NaCl, pH 7.5) in the absence of crowding agent. (**D**) HeLa cells were transiently transfected with siRNA against *PXN, FAK, ZYX*, or *GIT1*. Cell lysates were analyzed by immunoblotting against the indicated antibodies. (**E**) Full length PXN and HIC-5 recombinant proteins were separated on 10% SDS-PAGE gels, and stained with Coomassie Blue.

**Figure S4.** Characterization of PXN LLPS (related to Figure 5). (**A**) Purified PXN N-terminus (NT) and C-terminus (CT) were separated on 12% SDS-PAGE, and stained with Coomassie Blue. (**B**) Phase diagram of PXN at different protein concentrations and salt concentrations in the presence of 10% PEG 8000. (**C**) Representative images of PXN LLPS at salt concentrations ranging from 100 mM to 500 mM. PXN was labeled with 488 NHS ester. (**D**) Representative images at different time points showing the fusion event of PXN droplets. (**E**) The PXN droplets were analyzed by FRAP. The relative fluorescence intensity at different time points were quantified in (**F**). Data are shown as mean ± SD, n = 9. (**G**) MEFs stably expressing EGFP-PXN were analyzed by live cell imaging. Representative images captured the fusion of EGFP-PXN condensates. (**H**) FRAP analyses of EGFP-PXN condensates in the MEFs. The relative fluorescence intensity of the circled area was quantified and presented in (**I**). Data are shown as mean ± SD. n = 9. (**J**) *In vitro* co-phase separation of PXN with different FA proteins. PXN was labeled with 488 NHS ester and the other adhesion protein were labeled with Cy3 NHS ester. The reactions were in 150 mM NaCl, pH 7.5 in the presence of 10% PEG 8000. (**K**) Recombinant proteins of PXN WT, ΔIDR1, and ΔIDR2 were separated on 8% SDS-PAGE, and stained with Coomassie Blue. (**L**) *In vitro* pull-down assays using the indicated purified proteins.

**Figure S5.** Specific and non-specific molecular interactions cooperatively regulate FA assembly and integrin signaling (related to Figure 6). (**A**) Purified PXN WT and ΔLD2/4 proteins were run on SDS-PAGE gel, stained with Coomassie Blue. (**B**) Lysates prepared from *PXN*^-/-^ HeLa cells expressing empty vector (EV), PXN-Cry2 WT or ΔLD2/4 were analyzed by immunoblotting with the indicated antibodies. (**C**) Top, schematic illustration of PXN-Cry2 WT and ΔLD2/4. Bottom, representative images of *PXN*^-/-^ HeLa cells expressing PXN-Cry2 WT or ΔLD2/4 stimulated with blue light for 10 min. Number of Opto-PXN droplets was quantified in (**D**). ***p < 0.001 by unpaired Student t-test. (**E**) Representative images of *PXN*^-/-^ HeLa cells expressing PXN-Cry2 WT or ΔLD2/4 immunostained with antibodies against phospho-FAK (Tyr397), phospho-PXN (Tyr31 and Tyr118). (**F**) Representative images of *PXN*^-/-^ HeLa cells expressing PXN-Cry2 ΔLD2/4 immunostained with the indicated antibodies. (**G**, **H**) *PXN*^-/-^ HeLa cells reconstituted with empty vector (EV), PXN WT, or ΔLD2/4 were subjected to transwell assay. Representative images of cells stained with crystal violet were shown in (**G**) from 3 independent experiments. Quantification of cells migrated to the lower chambers was show in (**H**). Data are presented as mean ± SEM. ***p < 0.001 by one-way ANOVA.

## References

Alberti, S., A. Gladfelter, and T. Mittag. 2019. Considerations and Challenges in Studying Liquid-Liquid Phase Separation and Biomolecular Condensates. Cell. 176:419–434 https://doi.org/10.1016/j.cell.2018.12.035.

Alberti, S., and A.A. Hyman. 2021. Biomolecular condensates at the nexus of cellular stress, protein aggregation disease and ageing. Nat Rev Mol Cell Biol. 22:196–213 https://doi.org/10.1038/s41580-020-00326-6.

Alpha, K.M., W. Xu, and C.E. Turner. 2020. Paxillin family of focal adhesion adaptor proteins and regulation of cancer cell invasion. Int Rev Cell Mol Biol. 355:1–52 https://doi.org/10.1016/bs.ircmb.2020.05.003.

Bracha, D., M.T. Walls, and C.P. Brangwynne. 2019. Probing and engineering liquid-phase organelles. Nat Biotechnol. 37:1435–1445 https://doi.org/10.1038/s41587-019-0341-6.

Bracha, D., M.T. Walls, M.T. Wei, L. Zhu, M. Kurian, J.L. Avalos, J.E. Toettcher, and C.P. Brangwynne. 2018. Mapping Local and Global Liquid Phase Behavior in Living Cells Using Photo-Oligomerizable Seeds. Cell. 175:1467–1480 e1413 https://doi.org/10.1016/j.cell.2018.10.048.

Brangwynne, C.P., C.R. Eckmann, D.S. Courson, A. Rybarska, C. Hoege, J. Gharakhani, F. Julicher, and A.A. Hyman. 2009. Germline P granules are liquid droplets that localize by controlled dissolution/condensation. Science. 324:1729–1732 https://doi.org/10.1126/science.1172046.

Brown, M.C., J.A. Perrotta, and C.E. Turner. 1996. Identification of LIM3 as the principal determinant of paxillin focal adhesion localization and characterization of a novel motif on paxillin directing vinculin and focal adhesion kinase binding. J Cell Biol. 135:1109–1123 https://doi.org/10.1083/jcb.135.4.1109.

Case, L.B., M. De Pasquale, L. Henry, and M.K. Rosen. 2022. Synergistic phase separation of two pathways promotes integrin clustering and nascent adhesion formation. Elife. 11 https://doi.org/10.7554/eLife.72588.

Cirillo, L., A. Cieren, S. Barbieri, A. Khong, F. Schwager, R. Parker, and M. Gotta. 2020. UBAP2L Forms Distinct Cores that Act in Nucleating Stress Granules Upstream of G3BP1. Curr Biol. 30:698–707 e696 https://doi.org/10.1016/j.cub.2019.12.020.

Deakin, N.O., and C.E. Turner. 2008. Paxillin comes of age. J Cell Sci. 121:2435–2444 https://doi.org/10.1242/jcs.018044.

Deakin, N.O., and C.E. Turner. 2011. Distinct roles for paxillin and Hic-5 in regulating breast cancer cell morphology, invasion, and metastasis. Mol Biol Cell. 22:327–341 https://doi.org/10.1091/mbc.E10-09-0790.

Feng, Z., B. Jia, and M. Zhang. 2021. Liquid-Liquid Phase Separation in Biology: Specific Stoichiometric Molecular Interactions vs Promiscuous Interactions Mediated by Disordered Sequences. Biochemistry. 60:2397–2406 https://doi.org/10.1021/acs.biochem.1c00376.

Garabedian, M.V., W. Wang, J.B. Dabdoub, M. Tong, R.M. Caldwell, W. Benman, B.S. Schuster, A. Deiters, and M.C. Good. 2021. Designer membraneless organelles sequester native factors for control of cell behavior. Nat Chem Biol. 17:998–1007 https://doi.org/10.1038/s41589-021-00840-4.

Geiger, B., and K.M. Yamada. 2011. Molecular architecture and function of matrix adhesions. Cold Spring Harb Perspect Biol. 3 https://doi.org/10.1101/cshperspect.a005033.

Hagel, M., E.L. George, A. Kim, R. Tamimi, S.L. Opitz, C.E. Turner, A. Imamoto, and S.M. Thomas. 2002. The adaptor protein paxillin is essential for normal development in the mouse and is a critical transducer of fibronectin signaling. Mol Cell Biol. 22:901–915 https://doi.org/10.1128/MCB.22.3.901-915.2002.

Horton, E.R., A. Byron, J.A. Askari, D.H.J. Ng, A. Millon-Fremillon, J. Robertson, E.J. Koper, N.R. Paul, S. Warwood, D. Knight, J.D. Humphries, and M.J. Humphries. 2015. Definition of a consensus integrin adhesome and its dynamics during adhesion complex assembly and disassembly. Nat Cell Biol. 17:1577–1587 https://doi.org/10.1038/ncb3257.

Kanchanawong, P., G. Shtengel, A.M. Pasapera, E.B. Ramko, M.W. Davidson, H.F. Hess, and C.M. Waterman. 2010. Nanoscale architecture of integrin-based cell adhesions. Nature. 468:580–584 https://doi.org/10.1038/nature09621.

Kuo, J.C., X. Han, C.T. Hsiao, J.R. Yates, 3rd, and C.M. Waterman. 2011. Analysis of the myosin-II-responsive focal adhesion proteome reveals a role for beta-Pix in negative regulation of focal adhesion maturation. Nat Cell Biol. 13:383–393 https://doi.org/10.1038/ncb2216.

Legate, K.R., S.A. Wickstrom, and R. Fassler. 2009. Genetic and cell biological analysis of integrin outside-in signaling. Genes Dev. 23:397–418 https://doi.org/10.1101/gad.1758709.

Li, Y., T Zhang, H. Li, H. Yang, R. Lin, K. Sun, L. Wang, J. Zhang, Z. Wei, and C. Yu. 2020. Kindlin2-mediated phase separation underlies integrin adhesion formation. bioRxiv:2020.2007.2010.197400 https://doi.org/10.1101/2020.07.10.197400.

Li, Y.R., O.D. King, J. Shorter, and A.D. Gitler. 2013. Stress granules as crucibles of ALS pathogenesis. J Cell Biol. 201:361–372 https://doi.org/10.1083/jcb.201302044.

Mathieu, C., R.V. Pappu, and J.P. Taylor. 2020. Beyond aggregation: Pathological phase transitions in neurodegenerative disease. Science. 370:56–60 https://doi.org/10.1126/science.abb8032.

Mitra, S.K., D.A. Hanson, and D.D. Schlaepfer. 2005. Focal adhesion kinase: in command and control of cell motility. Nat Rev Mol Cell Biol. 6:56–68 https://doi.org/10.1038/nrm1549.

Parsons, J.T., A.R. Horwitz, and M.A. Schwartz. 2010. Cell adhesion: integrating cytoskeletal dynamics and cellular tension. Nat Rev Mol Cell Biol. 11:633–643 https://doi.org/10.1038/nrm2957.

Robertson, J., G. Jacquemet, A. Byron, M.C. Jones, S. Warwood, J.N. Selley, D. Knight, J.D. Humphries, and M.J. Humphries. 2015. Defining the phospho-adhesome through the phosphoproteomic analysis of integrin signalling. Nat Commun. 6:6265 https://doi.org/10.1038/ncomms7265.

Schuster, B.S., E.H. Reed, R. Parthasarathy, C.N. Jahnke, R.M. Caldwell, J.G. Bermudez, H. Ramage, M.C. Good, and D.A. Hammer. 2018. Controllable protein phase separation and modular recruitment to form responsive membraneless organelles. Nat Commun. 9:2985 https://doi.org/10.1038/s41467-018-05403-1.

Shin, Y., J. Berry, N. Pannucci, M.P. Haataja, J.E. Toettcher, and C.P. Brangwynne. 2017. Spatiotemporal Control of Intracellular Phase Transitions Using Light-Activated optoDroplets. Cell. 168:159–171 e114 https://doi.org/10.1016/j.cell.2016.11.054.

Shin, Y., and C.P. Brangwynne. 2017. Liquid phase condensation in cell physiology and disease. Science. 357 https://doi.org/10.1126/science.aaf4382.

Theodosiou, M., M. Widmaier, R.T. Bottcher, E. Rognoni, M. Veelders, M. Bharadwaj, A. Lambacher, K. Austen, D.J. Muller, R. Zent, and R. Fassler. 2016. Kindlin-2 cooperates with talin to activate integrins and induces cell spreading by directly binding paxillin. Elife. 5:e10130 https://doi.org/10.7554/eLife.10130.

Thomas, S.M., M. Hagel, and C.E. Turner. 1999. Characterization of a focal adhesion protein, Hic-5, that shares extensive homology with paxillin. J Cell Sci. 112 (Pt 2):181–190.

Turner, C.E., M.C. Brown, J.A. Perrotta, M.C. Riedy, S.N. Nikolopoulos, A.R. McDonald, S. Bagrodia, S. Thomas, and P.S. Leventhal. 1999. Paxillin LD4 motif binds PAK and PIX through a novel 95-kD ankyrin repeat, ARF-GAP protein: A role in cytoskeletal remodeling. J Cell Biol. 145:851–863 https://doi.org/10.1083/jcb.145.4.851.

Wang, Y., C. Zhang, W. Yang, S. Shao, X. Xu, Y. Sun, P. Li, L. Liang, and C. Wu. 2021. LIMD1 phase separation contributes to cellular mechanics and durotaxis by regulating focal adhesion dynamics in response to force. Dev Cell. 56:1313–1325 e1317 https://doi.org/10.1016/j.devcel.2021.04.002.

Winograd-Katz, S.E., R. Fassler, B. Geiger, and K.R. Legate. 2014. The integrin adhesome: from genes and proteins to human disease. Nat Rev Mol Cell Biol. 15:273–288 https://doi.org/10.1038/nrm3769.

Yang, P., C. Mathieu, R.M. Kolaitis, P. Zhang, J. Messing, U. Yurtsever, Z. Yang, J. Wu, Y. Li, Q. Pan, J. Yu, E.W. Martin, T. Mittag, H.J. Kim, and J.P. Taylor. 2020. G3BP1 Is a Tunable Switch that Triggers Phase Separation to Assemble Stress Granules. Cell. 181:325–345 e328 https://doi.org/10.1016/j.cell.2020.03.046.

Zacharchenko, T., X. Qian, B.T. Goult, D. Jethwa, T.B. Almeida, C. Ballestrem, D.R. Critchley, D.R. Lowy, and I.L. Barsukov. 2016. LD Motif Recognition by Talin: Structure of the Talin-DLC1 Complex. Structure. 24:1130–1141 https://doi.org/10.1016/j.str.2016.04.016.

Zhang, P., B. Fan, P. Yang, J. Temirov, J. Messing, H.J. Kim, and J.P. Taylor. 2019. Chronic optogenetic induction of stress granules is cytotoxic and reveals the evolution of ALS-FTD pathology. Elife. 8 https://doi.org/10.7554/eLife.39578.

Zhao, E.M., N. Suek, M.Z. Wilson, E. Dine, N.L. Pannucci, Z. Gitai, J.L. Avalos, and J.E. Toettcher. 2019. Light-based control of metabolic flux through assembly of synthetic organelles. Nat Chem Biol. 15:589–597 https://doi.org/10.1038/s41589-019-0284-8.

Zhu, J., Q. Zhou, Y. Xia, L. Lin, J. Li, M. Peng, R. Zhang, and M. Zhang. 2020. GIT/PIX Condensates Are Modular and Ideal for Distinct Compartmentalized Cell Signaling. Mol Cell. 79:782–796 e786 https://doi.org/10.1016/j.molcel.2020.07.004.

